# Division of labor between YAP and TAZ in non-small cell lung cancer

**DOI:** 10.1101/2020.01.12.894733

**Authors:** Michal Shreberk-Shaked, Bareket Dassa, Sanju Sinha, Silvia Di Agostino, Ido Azuri, Yael Aylon, Giovanni Blandino, Eytan Ruppin, Moshe Oren

## Abstract

Lung cancer is the leading cause of cancer-related deaths worldwide. The paralogous transcriptional cofactors Yes-associated protein (YAP) and transcriptional co-activator with PDZ-binding motif (TAZ, also called WWTR1), the main downstream effectors of the Hippo signal transduction pathway, are emerging as pivotal determinants of malignancy in lung cancer. Traditionally, studies have tended to consider YAP and TAZ as functionally redundant transcriptional cofactors, with similar biological impact. However, there is growing evidence that each of them also possesses distinct attributes. Here, we sought to systematically characterize the division of labor between YAP and TAZ in non-small cell lung cancer (NSCLC), the most common histological subtype of lung cancer. Employing representative NSCLC cell lines, as well as patient-derived data, we show that the two paralogs orchestrate non-overlapping transcription programs in this cancer type: whereas YAP preferentially regulates gene sets associated with cell division and cell cycle progression, TAZ preferentially regulates genes associated with extracellular matrix organization. Concordantly, depletion of YAP, but not TAZ, leads to growth arrest, while YAP overexpression promotes cell proliferation. Likewise, depletion of TAZ, but not YAP, compromises cell migration, whereas TAZ overexpression enhances migration. Importantly, the differential effects of YAP vs TAZ on key cellular processes are also associated with differential response to anti-cancer therapies. Uncovering the different activities and downstream effects of YAP and TAZ may thus facilitate better stratification of lung cancer patients for anti-cancer therapies.

## Introduction

Lung cancer is the leading cause of cancer-related deaths worldwide. Despite the significant progress that has been made toward understanding the causes of lung cancer, the 5-year survival is still lower than 15% [1]. Therefore, there is an urgent need for better understanding of the molecular mechanisms that drive lung cancer, which may eventually lead to more effective treatments.

The paralogous transcriptional cofactors Yes-associated protein (YAP) and transcriptional co-activator with PDZ-binding motif (TAZ, also called WWTR1), the main downstream effectors of the Hippo signal transduction pathway, are emerging as pivotal determinants of malignancy in lung cancer [2–7]. However, Hippo pathway genes, including *YAP/TAZ*, are rarely mutated in tumors, with only a few exceptions [8, 9], and thus YAP/TAZ dysregulation in cancer is likely to be driven by other mechanisms.

One of the open questions in the YAP/TAZ field is the extent to which these paralogs are functionally redundant. Although YAP and TAZ possess only about 57% amino acid sequence similarity (Supplementary Fig. S1), they are often regarded as functionally redundant, with similar biological impact [5, 10–13]. Indeed, both proteins play key roles in diverse biological processes, including tissue homeostasis, development, organ growth, regeneration, stem cell regulation and mechanotransduction [14–19]. Yet, a variety of structural and physiological features suggest that they may differ in their mode of regulation and their downstream activities [14, 20–22]. Notably, evidence for their different functions in lung cancer is also emerging [23, 24]. For example, YAP, but not TAZ, was reported to inhibit squamous trans-differentiation of lung adenocarcinoma cells [23]. Furthermore, Zhou and colleagues [4] demonstrated that TAZ protein levels are higher in lung cancer cell lines relative to normal lung epithelial cells, while YAP levels are more or less comparable in cancerous and non-cancerous lung cells. In addition, it was recently shown that lung cancer-derived cells depleted of *TAZ*, but not *YAP*, are more sensitive to bromodomain and extraterminal (BET) domain protein inhibitors [24]. Nevertheless, the extent of non-redundancy between YAP and TAZ in lung cancer cells, as well as the functional impact of such non-redundancy, remains to be comprehensively determined.

In the present study, we sought to explore the division of labor between YAP and TAZ in non-small cell lung cancer (NSCLC), the most common histological subtype of lung cancer [1]. Employing representative NSCLC cell lines, we now show that the two paralogs orchestrate non-identical transcription programs in these cells, giving rise to distinct biological phenotypes. Specifically, YAP preferentially regulates cell division and cell cycle progression, whereas TAZ preferentially regulates extracellular matrix, cell adhesion and cell migration. These findings imply that YAP and TAZ have distinct yet complementary roles in lung cancer cells. Identification of differential activities and downstream effects of YAP and TAZ may enable more refined stratification of lung cancer tumors, and uncover cancer vulnerabilities that might be exploited toward more effective therapy.

## Materials and Methods

### Cell lines and treatments

Cells were maintained at 37°C with 5% CO_2_. H1299 cells were cultured in RPMI-1640 (Gibco, USA) supplemented with 10% fetal bovine serum (Biological Industries, Israel) and 1% penicillin-streptomycin antibiotics solution (Biological Industries). A549 cells were grown in DMEM (Biological Industries) supplemented with 10% heat-inactivated fetal bovine serum (Hyclone, USA) and 1% penicillin-streptomycin antibiotics solution (Biological Industries). mCherry-labeled H1299 cells [25] were a kind gift of Uri Alon (Weizmann Institute of Science, Israel). A549 were obtained from ATCC. Cell lines were authenticated by STR profiling and tested negative for mycoplasma contamination. Experiments using cell lines were performed up to 9-12 passages after thawing from frozen stocks.

Taxol (Paclitaxel) was purchased from Sigma, USA (T7402) and was used at a final concentration of 1µM for 42 hours (for cell death assays).

### Western blot analysis

Western blot analysis was performed as described [26] with minor changes. Briefly, cells were washed with PBS, collected and lysed with NP-40 lysis buffer supplemented with phosphatase inhibitor cocktails 2 and 3 (Sigma) and protease inhibitor mix (Sigma). Lysates were centrifuged, and the supernatant was used to estimate protein concentration. Protein sample buffer was added, and samples were boiled for 5 minutes and loaded onto SDS-polyacrylamide (10%) gels for electrophoresis. Proteins were transferred onto nitrocellulose membranes, followed by 30 minutes blocking in 5% milk in Tris Buffered Saline with Tween (TBS-T). The membranes were incubated with primary antibodies overnight at 4°C, washed 3 times with TBS-T, and reacted for 45 minutes with horseradish-peroxidase (HRP)-conjugated IgG, followed by 3 washes with TBS-T. The proteins were visualized using an enhanced chemiluminescence (ECL) detection kit (Amersham, UK). Imaging and quantification were performed using a ChemiDoc MP imaging system (BioRad, USA) with the Image Lab 4.1 program (BioRad).

The following antibodies were used: GAPDH (Millipore Cat# MAB374, RRID:AB_2107445), YAP/TAZ (Cell Signaling Technology Cat# 8418, RRID:AB_10950494) and YAP (Santa Cruz Biotechnology Cat# sc-376830, RRID:AB_2750899). Conjugated anti-mouse or anti-rabbit secondary antibodies were from Jackson ImmunoResearch (USA).

### siRNA and plasmid transfections

For siRNA-mediated knockdown, the indicated SMARTpools or single oligonucleotides (Dharmacon, USA; see Table S2) were used with the Dharmafect#1 transfection reagent (Dharmacon) according to the manufacturer`s instructions, at a final concentration of 30nM. For siPLK1 knockdown we used 10nM PLK1 siRNA and 20nM siControl. The medium containing oligonucleotides and reagents was replaced after six hours. Plasmid transfection was done using jetPEI DNA transfection reagent (Polyplus Transfection, France), according to the manufacturer`s instructions. The final DNA amount was 10μg per 10cm dish, and the medium containing plasmids and reagents was replaced after five hours. pcDNA3-Flag-YAP and pcDNA3-Flag-TAZ were a generous gift of Yosef Shaul (Weizmann Institute of Science, Israel).

### RNA extraction, reverse transcription and qRT-PCR

Total RNA was isolated using a NucleoSpin RNA kit (Macherey Nagel, Germany). Reverse transcription and qPCR were performed as described in [27]. Values were normalized to either *HPRT* or *GAPDH*. Primers are listed in Table S2.

### RNA sequencing

H1299 cells were transfected with siControl, siYAP or siTAZ SMARTpool oligonucleotides for 48 hours and serum-starved for the last 18 hours. RNA was extracted and subjected to library preparation and RNA sequencing. Briefly, the polyA fraction was purified from 500ng of total RNA, followed by fragmentation and generation of double-stranded cDNA, Agencourt Ampure XP beads cleanup (Beckman Coulter, USA), end repair, A base addition, adapter ligation and PCR amplification. Libraries were quantified by Qubit (Thermo fisher scientific, USA) and TapeStation (Agilent, USA). Sequencing was done on Illumina HiSeq 2500 (Illumina, USA; single read sequencing).

### Transcriptomic analysis

RNA-sequencing analysis was done using the UTAP transcriptome analysis pipeline [28]. In short, reads were trimmed using cutadapt [29] and mapped to genome GRCh38 (Gencode) using STAR v2.4.2a [30] with default parameters. Genes having minimum 5 reads in at least one sample were considered. Normalization of the counts and detection of differential expression was performed using DESeq2 [31] with the betaPrior, cooksCutoff and independent filtering parameters set to False. Raw p-values were adjusted for multiple testing, using the procedure of Benjamini and Hochberg.

### Bioinformatic analysis

Visualization of gene expression heatmaps was done using Partek Genomics Suite 7.0 (Partek Inc., USA), using log normalized values (rld), with row standardization. Volcano plots were generated using Matlab.

### Gene ontology analysis

Gene Ontology (GO) analyses were performed with Metascape [32], using default parameters and “GO Biological Processes” functional set. GO terms with a q-value < 0.05 were considered significantly enriched.

### GSEA analysis

For GSEA analysis [33], genes were ranked according to fold change between the two described conditions. Comparison with indicated gene sets was done using GSEA pre-ranked tool. Enrichment of a specific dataset was considered significant when the false discovery (FDR) q-value was less than 0.05.

### Cell cycle profiling

H1299 or A549 cells were grown in 6cm plates and transfected with the indicated siRNAs or plasmids for 48 hours. Following serum starvation (0% serum) for 18 hours (unless stated otherwise), cells were harvested for cell cycle analysis using the Phase-Flow BrdU Cell Proliferation Kit (Biolegend, USA). Briefly, cells were incubated with BrdU for 75 minutes and labeled with an Alexa-647 conjugated anti-BrdU antibody. Total DNA was stained with DAPI. Then, 50,000 cells were collected and analyzed by multispectral imaging flow cytometry. The percentage of cells in each cell cycle phase was manually determined based on BrdU intensity and total DNA content, using FlowJo (Becton, Dickinson and Company, USA).

### Scratch assays

Cells were transfected with the indicated siRNAs or plasmids for 48 hours. When cells were approximately 90-100% confluent, a scratch was introduced with a 200µL pipette tip. Detached cells were washed off in serum-free medium. Gap closure was imaged at 0 and 24 hours with a Nikon eclipse Ti-E microscope at x4 magnification, capturing at least 4 fields for each condition. The migration distance was assessed manually using image J software (National Institutes of Health, USA). Gap closure was calculated using the following formula: (0hr gap width-24hr gap width)/0hr gap width.

### Cell death assays

Cell death was assessed using the CellTox Green Cytotoxicity Assay (Promega, USA) according to manufacturer’s instructions. Cells were seeded in a 96-well plate and transfected with indicated siRNAs or plasmids for 48 hours. Following incubation with either 1µM Taxol or DMSO for 42 hours, including serum starvation for the last 18 hours, CellTox reagent was added to the culture medium and the GFP fluorescence of each well was measured using an Infinite M200 microplate reader (Tecan, Switzerland). Three technical replicates were done for each condition. Cell death was calculated by subtracting the background signal and normalizing to DMSO-treated cells (control).

### Statistical Analysis

Independent biological replicates were performed and group comparisons were done as detailed in the figure legends. P-values below 0.05 were considered significant. Statistical significance between two experimental groups is indicated by asterisks.

### Synthetic lethality

We employed a recently published synthetic lethality (SL) pipeline, ISLE [34], to compute SL scores for all paralogous gene pairs (n=6353, derived from [35]). ISLE was performed on the expression profiles of 433 Lung adenocarcinoma (LUAD) human tumors from The Cancer Genome Atlas (TCGA) [36] and 87 LUAD cancer cell lines [37]. To compute the SL score for a given paralogous pair, the first step was to investigate the gene essentiality upon inactivation of its partner in LUAD cell lines. By definition, it is expected that gene A will be essential only when its SL partner gene B is inactive in a given cancer cell line [34]. Using a set of input genome-wide shRNA/sgRNA screens (as used in [34]), we tested whether the knockdown/knockout of one paralog is significantly more lethal when the other paralog is lowly expressed (bottom third quantile) vs highly expressed (top third quantile) across LUAD cell lines, via Wilcoxon rank-sum test (FDR corrected P<0.1). Second, underrepresented SL pairs were identified by quantifying the significance of simultaneous inactivation of the two paralogs together (under-expressed, via a hypergeometric test, FDR corrected P<0.1) in LUAD TCGA cohort. Third, we investigated the clinical relevance score, by performing a Cox multivariate regression testing whether the patients with an active SL interaction (both paralogs are inactive) have better survival than the rest of the patients, while controlling for cancer types, age, gender, race, tumor purity, genomic instability, and the effect of individual gene activation (FDR corrected P<0.1). To obtain a final SL score, we averaged the scores from these three steps, and scaled the scores from 0-1 (1 indicates high SL interaction). The paralogous pairs were ranked according to their final SL scores.

### YAP-differential and TAZ-differential drug sensitivity analysis

To identify YAP or TAZ distinct associations with drug response, we utilized drug sensitivity profiles (PRISM) of 4,518 drugs tested across 47 LUAD cancer cell lines [38]. We identified drugs that showed differential response between cell lines with high expression of *YAP* (top 33%) compared to those with low *YAP* expression (bottom 33%), and likewise for *TAZ*. P-value of the comparison was determined by Wilcoxon rank-sum test. Then, by using Drugbank [39] annotation, the targets for each of the YAP-differential or TAZ-differential drugs were defined. YAP-differential and TAZ-differential drugs, their targets and the p-values are listed in Table S4.

### Taxol sensitivity analysis

#### PRISM dataset

By utilizing single dose Taxol cell viability data across cancer cell lines (n=578) [38], we tested whether lines with high expression of YAP (top 33%, n=192) are differentially more sensitive to Taxol (Paclitaxel) compared to those with low YAP expression (bottom 33%, n=192), and likewise for TAZ. P-value of the comparison was determined by Wilcoxon rank-sum test.

#### GDSC dataset

Taxol sensitivity data was obtained from [40], in which sensitivity was measured by area under the dose-response curve in 136 lung cancer cell lines. The Spearman correlation coefficient of Taxol sensitivity with either YAP or TAZ log2-transformed protein levels (determined by RPPA in CCLE [41]) was calculated across these cell lines.

### TCGA YAP and TAZ correlation analysis

We queried the LUAD expression dataset of TCGA, comprising 20,167 expressed genes and 515 tumor samples (after cleaning and filtering the data). LUAD TCGA mRNA-seq RSEM normalized data was downloaded from http://gdac.broadinstitute.org/. Expression data (X) was transformed to X_transformed = log2(X + 1). The Pearson correlation coefficient with either *YAP* or *TAZ* mRNA level was calculated for each gene. Genes were sorted according to absolute R, and the top eight percent (1600 genes) were selected for further analysis. Non-overlapping genes between the YAP-correlated and TAZ-correlated lists (1276 genes) were subjected to GO annotation analysis.

## Results

### YAP and TAZ are associated with distinct transcriptional programs

YAP and TAZ act primarily as transcriptional cofactors [10, 13, 15]. To compare YAP and TAZ impact on the transcriptome of lung cancer-derived cells, we performed RNA-sequencing (RNA-seq) analysis following siRNA-mediated transient knockdown of either *YAP* (siYAP) or *TAZ* (siTAZ). As a model system we chose the NSCLC-derived H1299 cell line, commonly used in lung cancer research [42–44]. Knockdown validation is shown in Fig. 1A. As expected, a subset of common genes (82 in total) were impacted to a similar extent by partial depletion of either YAP or TAZ (Fig. 1B). However, expression of a larger number of genes was differentially affected in a paralog-specific manner. Thus, 204 genes were significantly downregulated or upregulated at least 2-fold upon YAP depletion, but were only mildly affected by TAZ depletion (Fig. 1B, C). Conversely, the expression of 324 other genes was significantly affected by TAZ depletion, but less so by YAP depletion (Fig. 1B, D). Hence, while both paralogs regulate to a similar extent a subset of common genes, expression of a substantially greater number of genes in these cells is differentially dependent on one of the paralogs as compared to the other one. For simplicity, the sets of genes preferentially regulated by YAP or TAZ will hereafter be referred to as YAP-regulated or TAZ-regulated genes, respectively.

**Figure 1:**
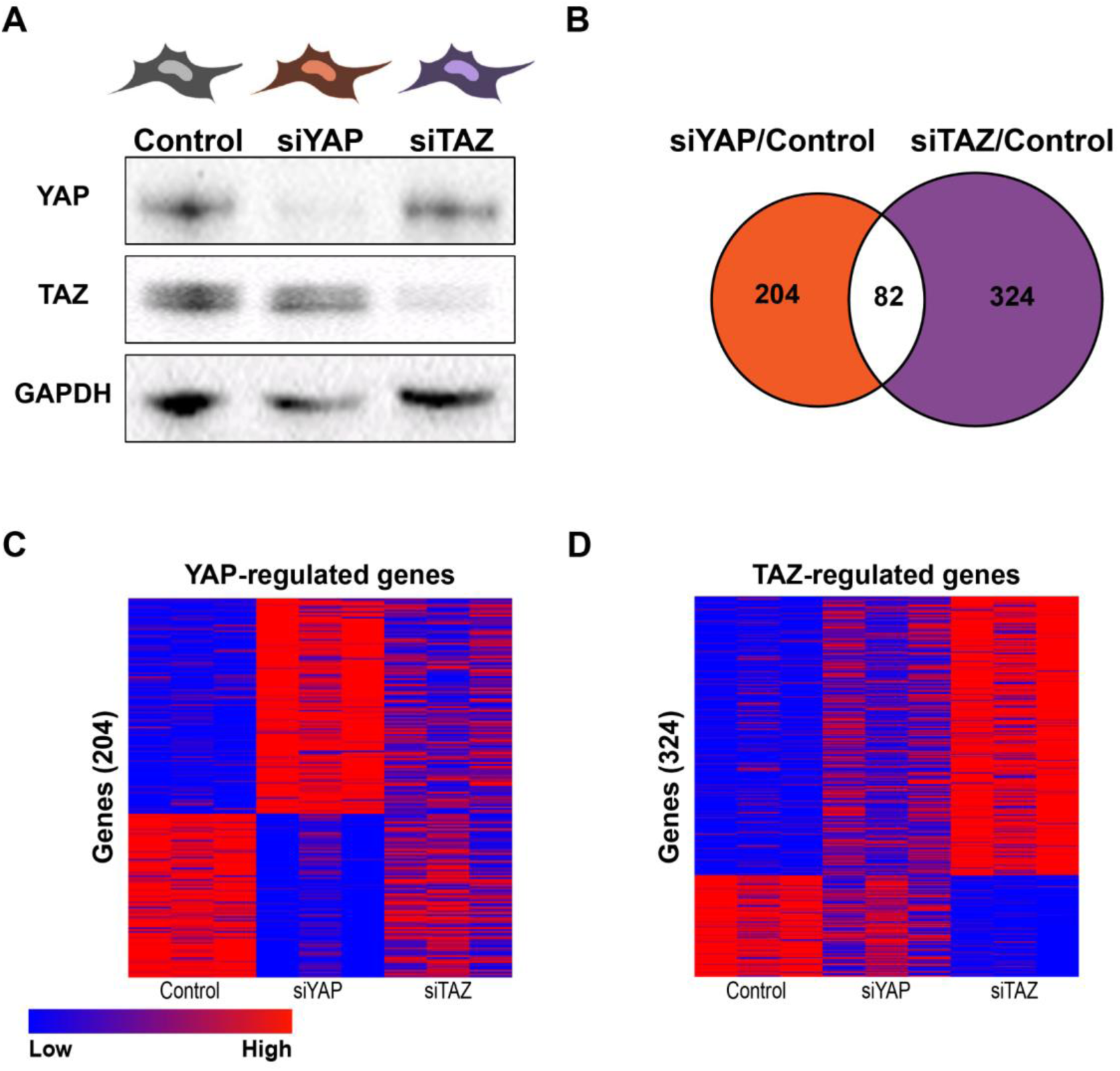
YAP and TAZ are associated with distinct transcriptional programs in H1299 cells. A. Human NSCLC-derived H1299 cells were transiently transfected with siRNA SMARTpools targeting either YAP (siYAP) or TAZ (siTAZ) or with control siRNA; RNA was extracted 48 hours later and subjected to RNA-seq analysis. Upper panel: Cartoon describing the approach taken to determine YAP and TAZ transcriptional programs. Lower panel: Representative immunoblot with the indicated antibodies. B. Venn diagram of the overlap between significantly differentially expressed genes upon transfection with either siYAP or siTAZ compared to control, from three biological repeats. Orange and purple indicate genes whose expression was selectively altered (either positively or negatively) by siYAP or siTAZ, respectively. Absolute fold change≥2, adjusted p-value ≤0.05. C and D. Heatmaps of gene-expression levels of the non-overlapping 204 siYAP significantly differentially expressed genes (C) and the 324 siTAZ significantly differentially expressed genes (D) depicted in (B).

### YAP preferentially regulates cell cycle progression while TAZ preferentially regulates cell migration

To identify key biological processes regulated by either YAP or TAZ in H1299 cells, we performed gene ontology analyses on the YAP-regulated or TAZ-regulated genes. Interestingly, whereas the genes preferentially regulated by YAP were strongly enriched for cell division and mitosis-related terms (Fig. 2A and Table S1), the top enriched terms for the TAZ-regulated genes were related to extracellular matrix (ECM) organization (Fig. 2B and Table S1). Notably, we observed that almost all the YAP-associated cell division genes were downregulated upon *YAP* silencing, implying that they are positively regulated by YAP (Fig. 2C). In contrast, the majority of ECM-related genes included in the TAZ-specific signature were upregulated upon *TAZ* silencing (Fig. 2D), indicating that their expression is repressed by TAZ in these cells.

**Figure 2:**
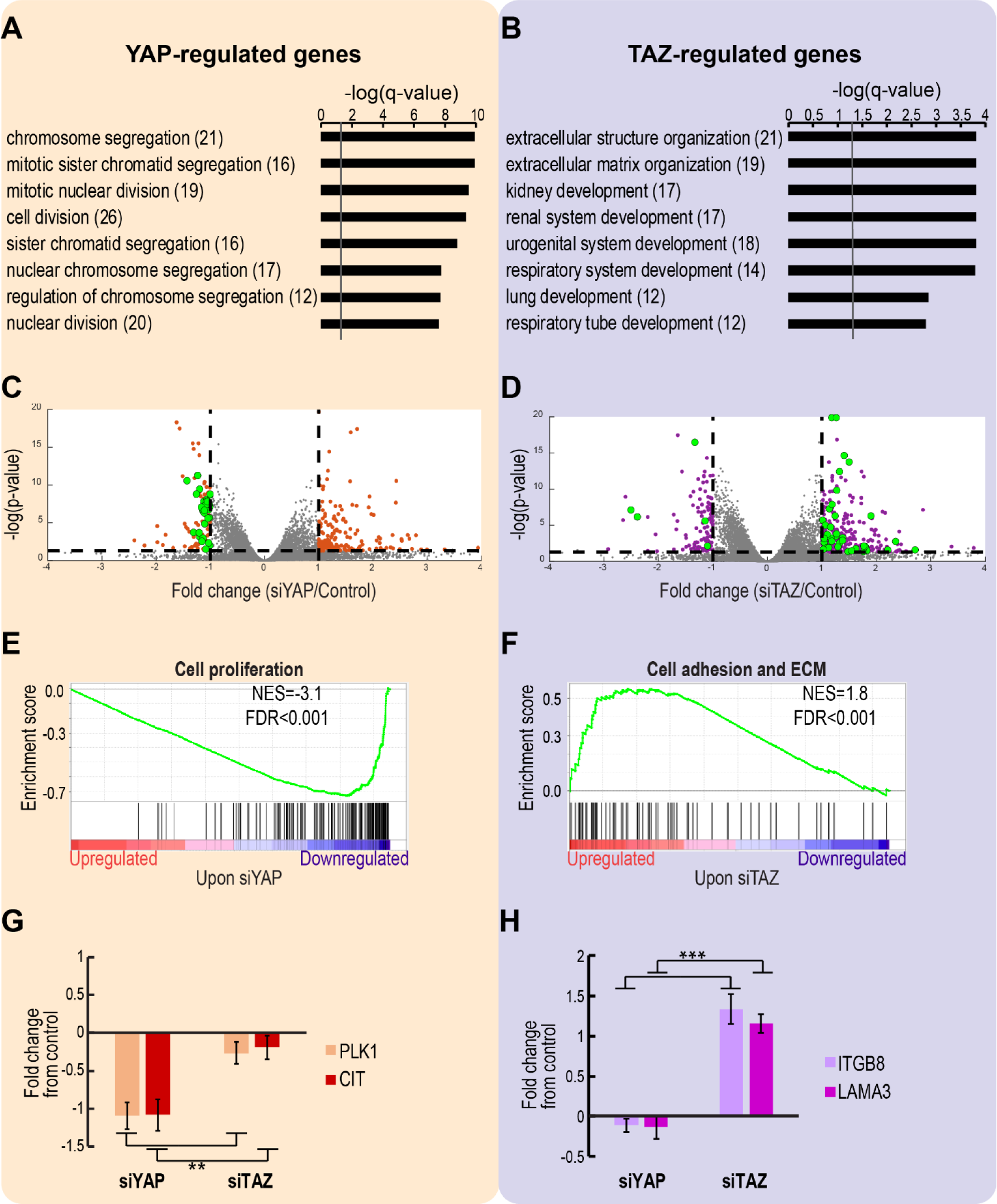
YAP-regulated genes are enriched for cell division terms, while TAZ-regulated genes are enriched for ECM and adhesion related terms. A and B. Eight top enriched GO Biological Processes terms of the YAP-regulated (A) and TAZ-regulated (B) genes, determined by Metascape. Q-value is –log10 transformed; gray line represents a q-value of 0.05. In brackets is the number of enriched genes in each term. C and D. Volcano plots of 18,977 genes detected in the RNA-seq analysis. Colored dots represent genes whose expression was significantly altered by either siYAP (orange) or siTAZ (purple), relative to control siRNA. Genes related to the top enriched GO terms for each siRNA (black columns in Fig. 2A, B) are marked in green. P-value represents corrected p-value and is −log10 transformed. Dashed lines represent p-value of 0.05 (bottom), and log2 fold change of 2 (right) and 0.5 (left). For extended plots, see Supplementary Fig. S2. E and F. GSEA enrichment plots for all the analyzed genes from the RNA-seq, ranked by fold change upon either siYAP or siTAZ relative to control, compared to gene sets associated with the pathways indicated above each plot. NES = Normalized enrichment score; FDR = False discovery rate. G and H. RT-qPCR analysis of representative cell proliferation-related (G) and ECM-related (H) genes. Data represent log2 mRNA expression (mean+SEM) normalized to HPRT and control transfected cells, from six independent biological repeats. **p<0.01; ***p<0.001 one-way ANOVA and Tukey’s post hoc test of the indicated comparisons.

Gene Set Enrichment Analysis (GSEA) on all informative genes ranked by their expression fold change (relative to control) upon either siYAP or siTAZ showed that the top downregulated genes upon *YAP* silencing were strongly enriched for cell proliferation [45] (Fig. 2E), whereas the genes upregulated upon *TAZ* silencing were enriched for adhesion and ECM [46] (Fig. 2F).

Selected genes from each subset were validated by RT-qPCR analysis. As shown in Fig. 2G, canonical cell cycle regulators, such as PLK1 and CIT [47–49] were indeed preferentially downregulated upon siYAP, but not siTAZ. Likewise, integrins and laminins, including ITGB8 and LAMA3, which affect cell adhesion and migration [50, 51], were significantly upregulated upon silencing of *TAZ*, but not *YAP* (Fig. 2H).

We next aimed to determine whether the differential transcriptomic effects of YAP and TAZ lead to different functional outcomes. We therefore examined the effects of depletion of YAP or TAZ on the proliferation and migration of two NSCLC-derived cell lines, H1299 and A549.

Depletion of either *PLK1* or *CIT* genes, preferentially regulated by YAP (Fig. 2G), has been reported to attenuate cell cycle progression in multiple cancer cell types [47, 49, 52], leading to depletion of the S-phase subpopulation and accumulation of cells in G1-phase. Indeed, as seen in Fig. 3A, B, silencing of *YAP*, but not *TAZ*, resulted in a prominent depletion of S-phase cells in both cell lines (siYAP/Control ratio = 0.4 in H1299 cells, and 0.3 in A549 cells) and elicited a mild increase in the G1 subpopulation, relative to control or siTAZ. Similarly, *PLK1* silencing mimicked the cell cycle effects of siYAP (Supplementary Fig. S3). Conversely, transient overexpression of YAP, but not TAZ (Supplementary Fig. S4), increased the S-phase fraction (Fig. 3C). Altogether, our observations imply that YAP, but not TAZ, acts as a positive regulator of cell cycle progression in these lung cancer cells.

**Figure 3:**
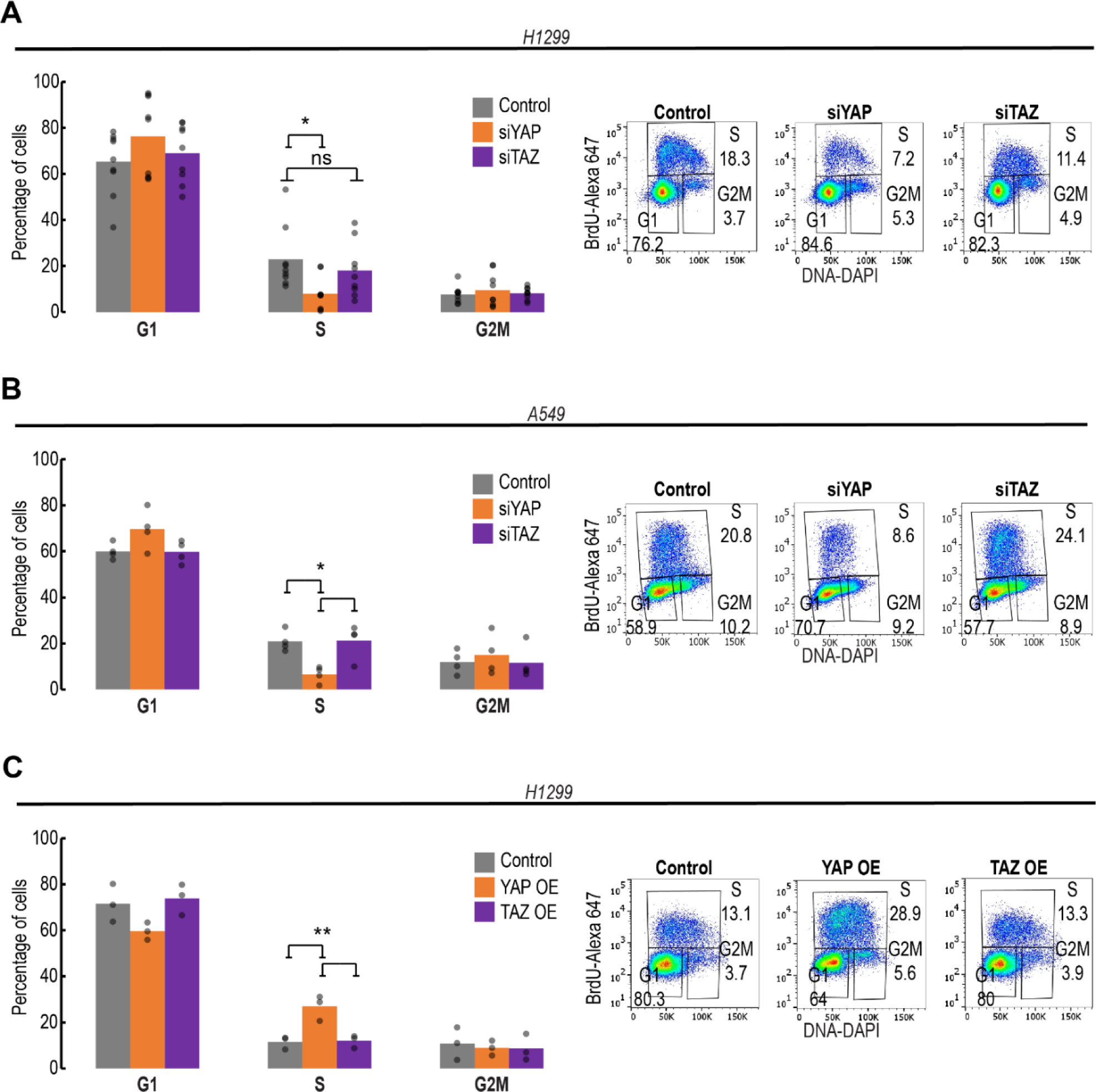
YAP, but not TAZ, preferentially promotes cell cycle progression in NSCLC-derived cells. A and B. Cell cycle profiling, by BrdU + DAPI analysis, of H1299 (n=9) and A549 (n=4) cell cultures transfected with the indicated siRNAs. C. BrdU + DAPI cell cycle profiling of H1299 cell cultures (n=3) transfected transiently with plasmids encoding either YAP-flag (YAP OE) or TAZ-flag (TAZ OE), or control plasmid. Left panels: average percentages of cells in each cell cycle phase; each dot represents an independent biological repeat. Right panels: representative FACS analysis images. ns = not significant; *p<0.05; **p<0.01 determined by one-way ANOVA and Tukey’s post hoc test of the indicated comparisons.

To assess the impact of YAP and TAZ on cell migration, we performed gap closure (“scratch”) assays. Excessive ECM production can impede cell migration, owing to aberrant cell adhesion [53]. Indeed, *TAZ* knockdown, which augmented the expression of adhesion and ECM-related genes (Fig. 2B,F), strongly attenuated cell migration in both H1299 and A549 cells, whereas silencing of *YAP* had almost no effect (Fig. 4A,B). Concordantly, transient overexpression of TAZ, but not YAP, led to a mild increase in cell migration (Fig. 4C). Silencing of *YAP* or *TAZ* by single siRNA oligonucleotides phenocopied the effects of the corresponding siRNA SMARTpools (Supplementary Fig. S5). Altogether, our observations imply that YAP and TAZ have non-overlapping roles in these lung cancer cells: whereas YAP preferentially promotes cell cycle progression, TAZ preferentially promotes cell migration.

**Figure 4:**
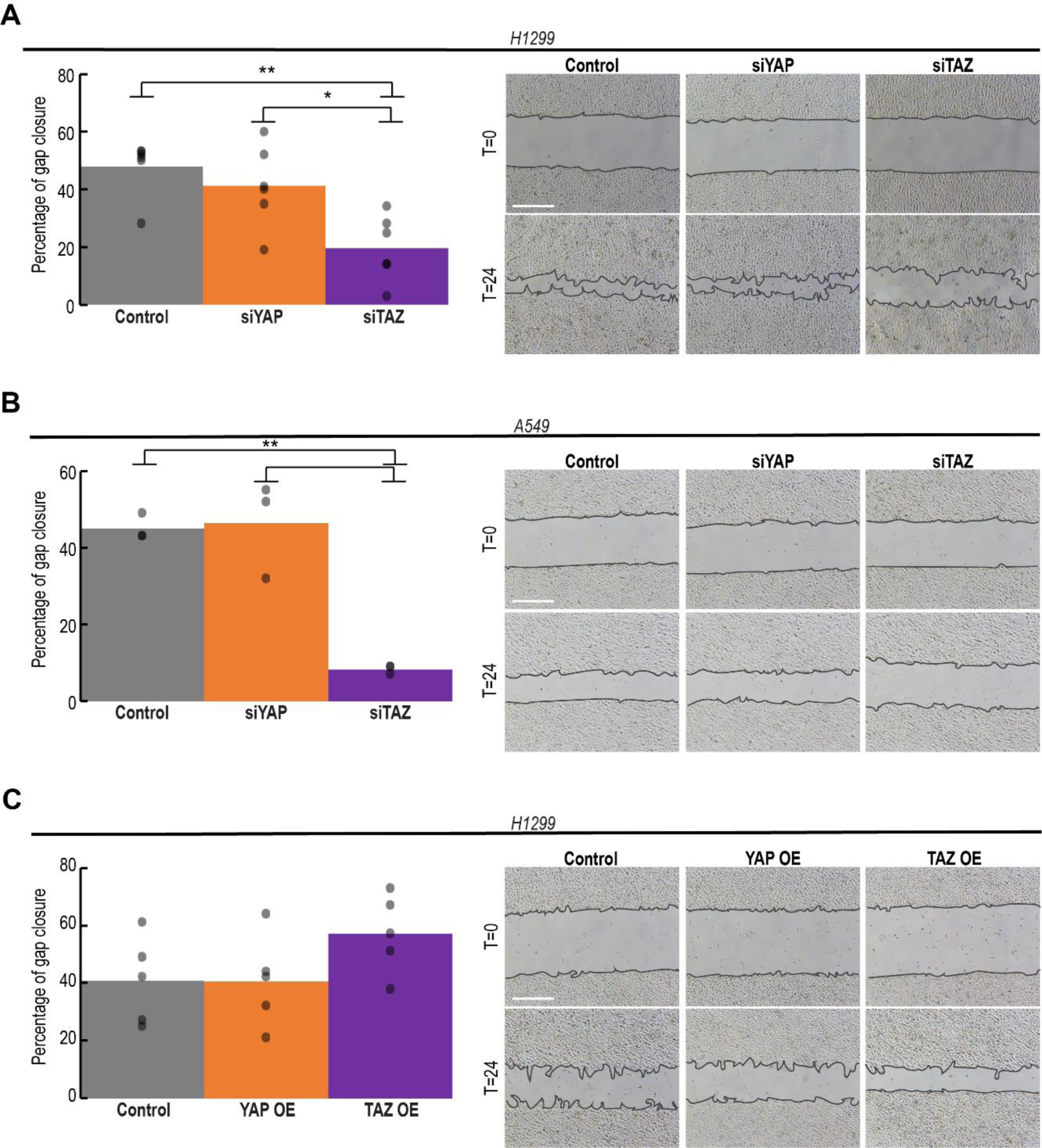
TAZ, but not YAP, preferentially promotes cell migration. A and B. Gap closure (“Scratch”) assays of H1299 (n=6) and A549 (n=3) cell cultures transfected with the indicated siRNAs. C. Gap closure assay of H1299 cell cultures (n=5) transfected transiently with plasmids encoding either YAP-flag (YAP OE) or TAZ-flag (TAZ OE), or control plasmid. Left panels: Average percentage of gap closure calculated from all biological repeats; each dot represents an independent biological repeat. Right panels: representative images of gap closure at T=0 and T=24 hours. *p<0.05; **p<0.01 determined by one-way ANOVA and Tukey’s post hoc test of the indicated comparisons. Scale bar = 500µM.

### YAP and TAZ display partial functional non-redundancy across lung cancer cell lines and tumors and are associated with distinct cancer-associated phenotypes in lung tumors

One broadly accepted way to study functional backup and redundancy between genes is by examining their extent of synthetic lethality (SL) [54]. SL implies a genetic interaction between two genes, whereby the individual inactivation of either gene results in a viable phenotype, while their combined inactivation reduces cell fitness (and in extreme cases, is lethal) [34]. Redundant paralogous pairs often display strong synthetic lethality, such that cells expressing intrinsically low levels of one paralog are highly dependent on retention of the other paralog for their survival [35]. Hence, the extent of SL informs on the extent of functional redundancy of a paralogous pair. The partially non-redundant transcriptional and functional effects of YAP and TAZ in lung cancer cells suggested that these paralogs might display only partial synthetic lethality. To test this prediction, we utilized a recently published SL identification computational pipeline [34] to compute SL scores of human paralogs, as previously defined in the literature (n=6353) [35], in lung adenocarcinoma (LUAD) cells and tumors (see Materials and Methods). Indeed, as shown in Fig. 5A, we found that YAP-TAZ have a fairly low degree of SL when compared to the ranking of all paralogous pairs (SL Score = 0.37, Rank= 5643). In contrast, ARID1A and ARID1B, known to be functionally redundant [55], were ranked much higher (SL Score= 0.95, Rank=8, Fig. 5A, green color). As negative controls, we paired the ARID1A-ARID1B paralogous pair with either YAP or TAZ (YAP-ARID1A, YAP-ARID1B, TAZ-ARID1A, TAZ-ARID1B); as expected, these pairings yielded very low SL scores (mean SL Score = 0.12) (Fig. 5A, red color). Thus, the SL analysis further confirms that YAP and TAZ are only mildly redundant in lung cancer cells.

**Figure 5:**
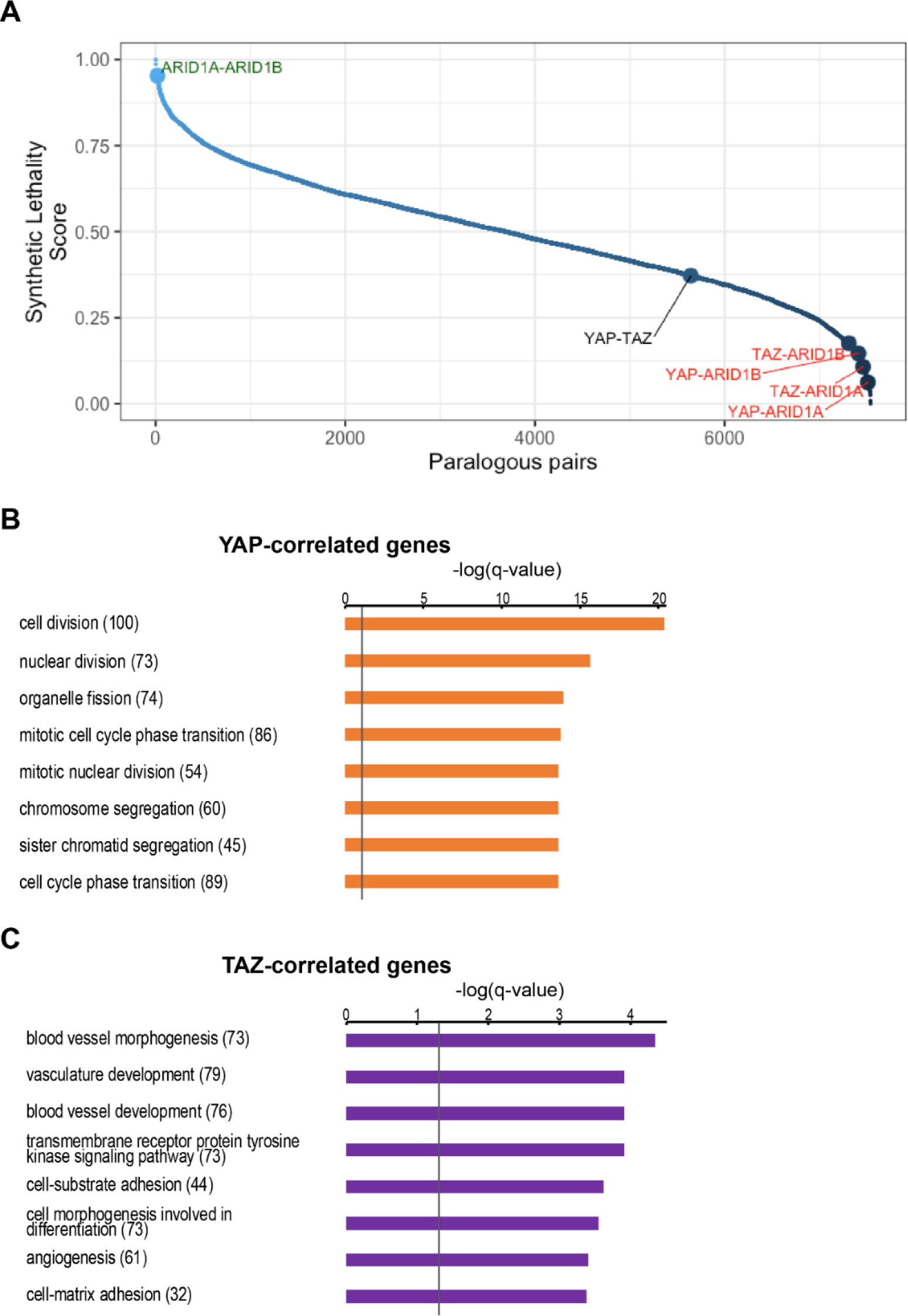
YAP and TAZ display partial non-redundancy and their expression is correlated with distinct biological processes in lung adenocarcinoma cell lines and tumors. A. Ranked synthetic lethality scores of all paralogous pairs, computed via the ISLE pipeline [34], in LUAD cell lines and tumors (see Materials and Methods). Higher score indicates higher synthetic lethality. Representative positive control and negative control pairs are shown in green and red, respectively. B and C. Gene ontology analysis (performed by Metascape) of the non-overlapping YAP-correlated and TAZ-correlated genes, regardless of directionality (absolute R) in lung adenocarcinoma tumors (TCGA LUAD). The top eight enriched GO Biological Processes terms are shown. Gray line represents q-value of 0.05. In brackets is the number of enriched genes in each term.

Next, we asked whether expression levels of *YAP* and *TAZ* in lung tumors are associated with cell cycle or ECM-related terms. Toward this aim, we queried the LUAD expression dataset of The Cancer Genome Atlas (TCGA). Utilizing an unbiased correlation-based method, the correlation coefficient with either *YAP* or *TAZ* mRNA was calculated for each gene. Interestingly, *YAP* and *TAZ* mRNA levels are only partially correlated (Supplementary Fig. S6). The top 1600 genes displaying the highest correlation coefficients (regardless of directionality), were further analyzed. Remarkably, only 324 of the 1600 genes overlapped between the YAP-correlated and TAZ-correlated gene sets. Importantly, gene ontology analysis of the non-overlapping genes (n=1276) showed that the YAP-correlated genes were enriched in cell division and cell cycle progression terms (Fig. 5B and Table S3), while TAZ-correlated genes were enriched in morphogenesis, development and adhesion related terms (Fig. 5B and Table S3); of note, morphogenesis is linked with adhesion related genes [56]. Hence, our *in vitro* observations, implying distinct roles for YAP and TAZ in lung cancer cell biology, also appear to hold for actual human cancer patients.

### Differential association of YAP vs TAZ with response to anti-cancer drugs

Direct pharmacological inhibition of YAP/TAZ activity is of growing interest, but still remains a clinical challenge [57]. Nevertheless, the distinct functions of YAP and TAZ suggest that the two paralogs may affect differently also the response of cancer cells to particular anti-cancer therapies. Indeed, using drug sensitivity data across 47 LUAD cell-lines (PRISM dataset [38]), we identified 142 drugs whose response profile differs preferentially between high vs low *YAP* mRNA levels, and 38 drugs whose response profile differs preferentially between high vs low *TAZ* mRNA levels (Table S4).

Taxol (Paclitaxel), a microtubule poison, is among the drugs used to treat NSCLC [58]. Interestingly, when inspecting the entire collection of cell lines comprised in the drug sensitivity dataset [38], we observed that cell viability upon Taxol treatment was negatively correlated with the expression levels of *YAP*, but not *TAZ* (Fig. 6A). A further analysis of the largest publicly available cancer cell line drug sensitivity collection (GDSC dataset [40]) confirmed that Taxol sensitivity, measured by area under the dose-response curve, was more significantly correlated with YAP protein levels (determined by RPPA) (R=0.22, p-value<0.02, Supplementary Fig. S7) than with TAZ protein levels (R=0.10, p-value = 0.23, Supplementary Fig. S7) in lung cancer cell lines.

**Figure 6:**
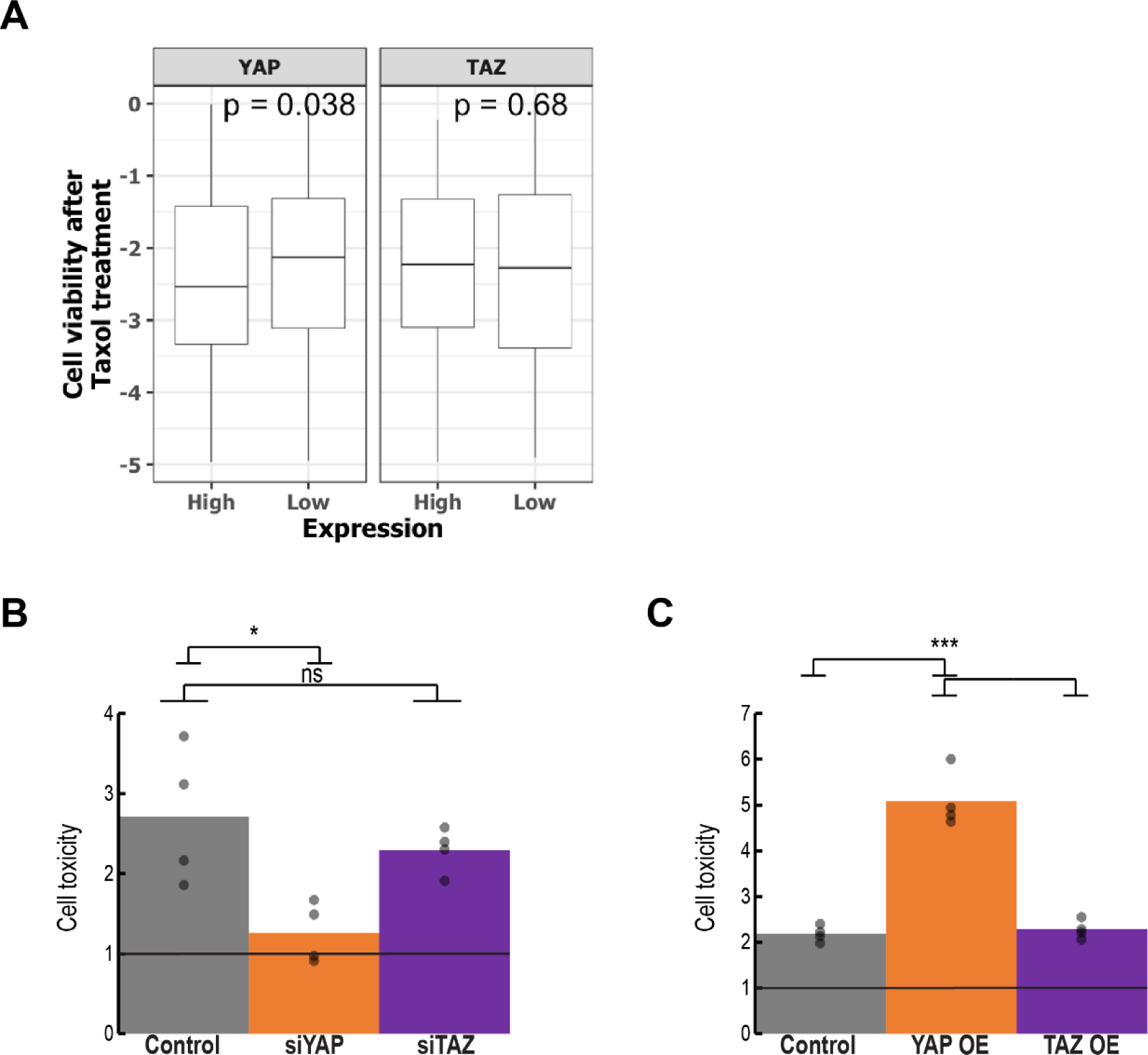
Taxol sensitivity is preferentially dependent on YAP. A. Box plot of Taxol cell viability scores across cancer cell lines (n=578) derived from PRISM [38], binned according to YAP or TAZ expression levels; lower cell viability implies greater sensitivity to Taxol. P = p-value determined by Wilcoxon rank-sum test. C and D. H1299 cells were transfected with the indicated siRNA SMARTpools (C) or expression plasmids (D), and 6 hours later were subjected to treatment with 1µM Taxol for an additional 42 hours and serum starvation for the last 18 hours. The percentage of dead cells was determined by the CellTox Green Cytotoxicity Assay (see Materials and Methods), and cell death was calculated relative to DMSO-treated cells. Average cell death calculated from four biological repeats; each dot represents an independent repeat. ns = not significant; *p<0.05; ***p<0.001, determined by one-way ANOVA and Tukey’s post hoc test of the indicated comparisons.

To compare directly the impact of YAP vs TAZ on Taxol sensitivity, we subjected H1299 cells to Taxol treatment after knockdown of either *YAP* or *TAZ*. As seen in Fig. 6B, *YAP* silencing indeed greatly compromised the ability of Taxol to trigger cell death, while *TAZ* silencing had only a marginal effect. Conversely, transient overexpression of *YAP* led to increased sensitivity to Taxol, which was not seen with *TAZ* overexpression (Fig. 6C). Thus, in agreement with their distinct biological roles in lung cancer cells, YAP and TAZ also affect differentially the sensitivity to specific anti-cancer agents.

## Discussion

In the present study, we compared the functional impact of partial depletion of YAP versus that of TAZ, in NSCLC-derived cells. We found that while both paralogs modulated to a similar extent a common subset of shared genes, distinctly larger subsets were preferentially affected in a paralog-specific manner. In agreement with the different compositions of these subsets, there was also a pronounced difference in biological impact: whereas YAP strongly affected cell cycle progression but not cell migration, TAZ modulated migration but not cell cycle progression. This differential association was apparent not only in cell lines, but also in data from human lung tumors. Thus, there exists a division of labor between the two paralogs in lung cancer cells and tumors. Moreover, this division of labor is associated with differential responses to anti-cancer drugs, and specifically to Taxol. For the majority of genes, this division is preferential rather than absolute, as many YAP-regulated genes were also mildly affected by TAZ depletion, and vice versa (Fig. 1C,D). This suggests that the relative contribution of YAP vs TAZ to the expression of such genes may also depend on the relative abundance of each paralog in a given cell. More broadly, a similar idea was recently suggested regarding protein-protein interactions: two paralogs may have overlapping potential binding partners, but the actual interaction profile of each paralog in a given cellular setting will be determined by its relative abundance and relative affinities for the different interaction partners [59]. Therefore, the division of labor between YAP and TAZ is expected to be context-dependent, and may differ greatly between different types of cancer and perhaps also between individual cases of the same cancer type. Interestingly, Sun *et al.* [60] have recently shown that while YAP and TAZ have overlapping functions in promoting the proliferation of non-differentiated myoblasts, they exert opposing effects when such cells are induced to undergo myogenic differentiation: whereas TAZ enhances differentiation, YAP actually inhibits it. Furthermore, Plouffe *et al.* [61] found that inactivation of *YAP* in human embryonic kidney cells had a greater effect on several cellular processes than inactivation of *TAZ*. Both of these studies [60, 61] support the notion that one paralog may be more dominant over the other in regulating specific biological functions.

Gene duplication events have occurred in many instances in the course of evolution, and have played a major role in shaping the genomes of different species, including humans. Paralogs originating from a gene duplication event can diverge and acquire changes that contribute to increased organismal diversity. Invertebrates have only a single *YAP/TAZ* orthologue; in *Drosophila* this orthologue, *Yorkie* (*Yki*), is required for both cell proliferation and cell migration in the course of multiple developmental processes [62–66]. In vertebrates, while both YAP and TAZ can regulate both cell proliferation and cell migration, they do so with different specificities, thereby enabling more refined regulation of complex biological processes and conditions. We propose that the gene duplication and the consequent division of labor between the YAP/TAZ paralogs might explain their retention throughout vertebrate evolution. Interestingly, *Yap* knockout mice are embryonic lethal [67], while *Taz* knockout mutants are partially viable [68, 69], further supporting their non-redundancy.

The molecular mechanisms underpinning the differential effects of YAP and TAZ in lung cancer cells still remain to be elucidated. Both YAP and TAZ do not bind directly to DNA, but rather are recruited to the chromatin indirectly by “piggybacking” on defined DNA-binding proteins, which recognize distinct sequences within the genomic DNA [10, 11]. Hence, it is plausible that YAP and TAZ may vary in their relative affinities for particular DNA binding proteins, which may lead to different transcriptional preferences, particularly when such DNA binding proteins are present in limited amount. Such relative affinities might be subject to further regulation, e.g. by post-translational modifications of YAP or TAZ, providing an additional layer of context-dependent diversity. Furthermore, it was recently demonstrated [70] that TAZ, but not YAP, can form nuclear condensates via liquid-liquid phase separation (LLPS) to compartmentalize its DNA binding co-factor TEAD4, the transcription co-activators BRD4 and MED1 and the transcription elongation factor CDK9 for activation of gene expression. Yet, YAP is capable of forming LLPS condensates under a different set of experimental conditions [71], further highlighting the context-dependent nature of the similarities/differences between the two paralogs.

Regardless of mechanisms, our findings suggest that tumorigenesis may take advantage of the division of labor between YAP and TAZ in order to orchestrate complementary oncogenic functions. These findings may enable a more refined stratification of lung cancer tumors, and perhaps reveal cancer vulnerabilities that might be exploited toward more effective therapies.

## Supplementary data

**Figure S1:**
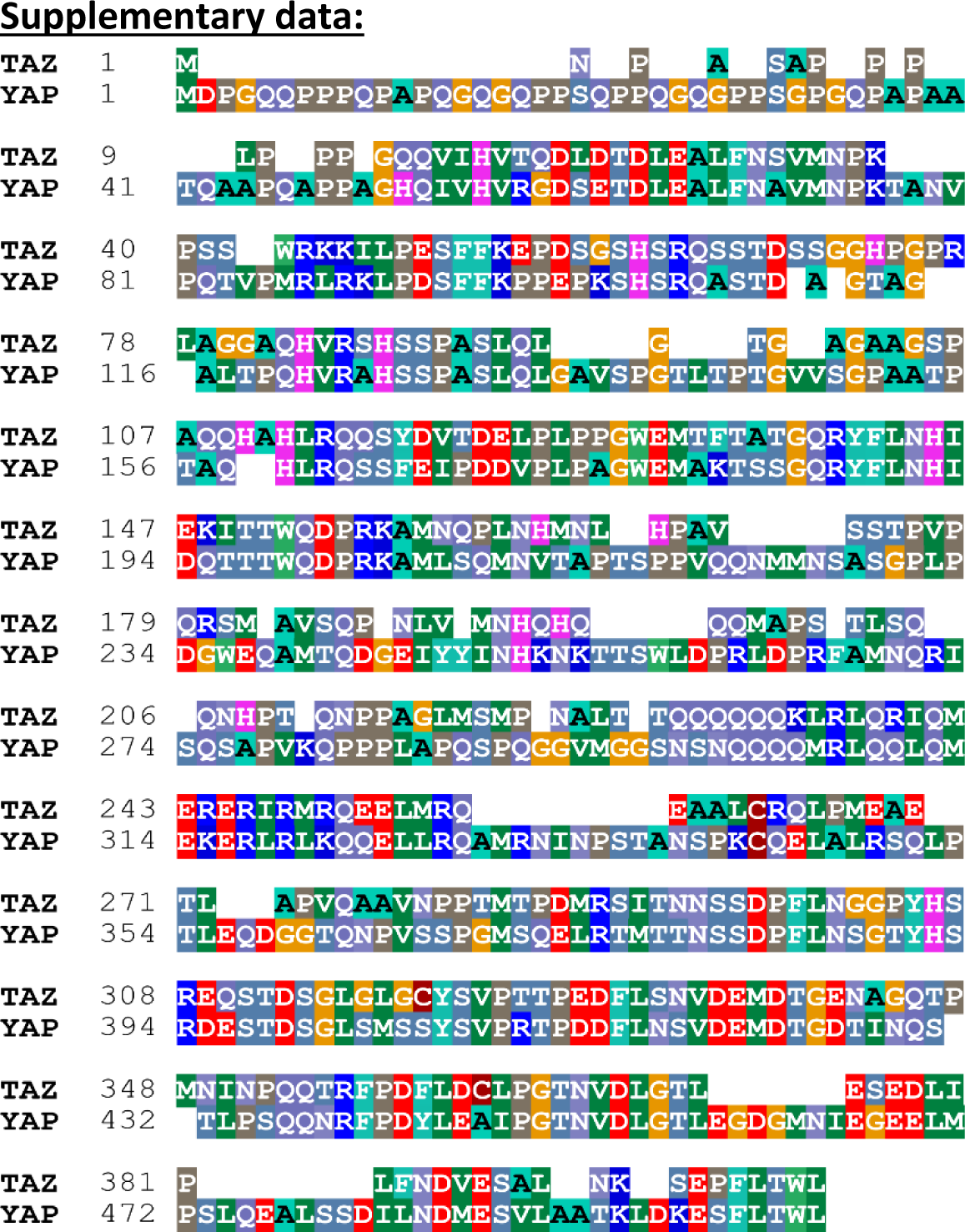
Comparison of YAP and TAZ protein sequences. Pairwise sequence alignment of the protein sequences of TAZ and YAP. Numbers at the left relate to amino acid positions. Color shading denotes different groups of amino acids. Alignment was generated by using BioEdit software.

**Figure S2.**
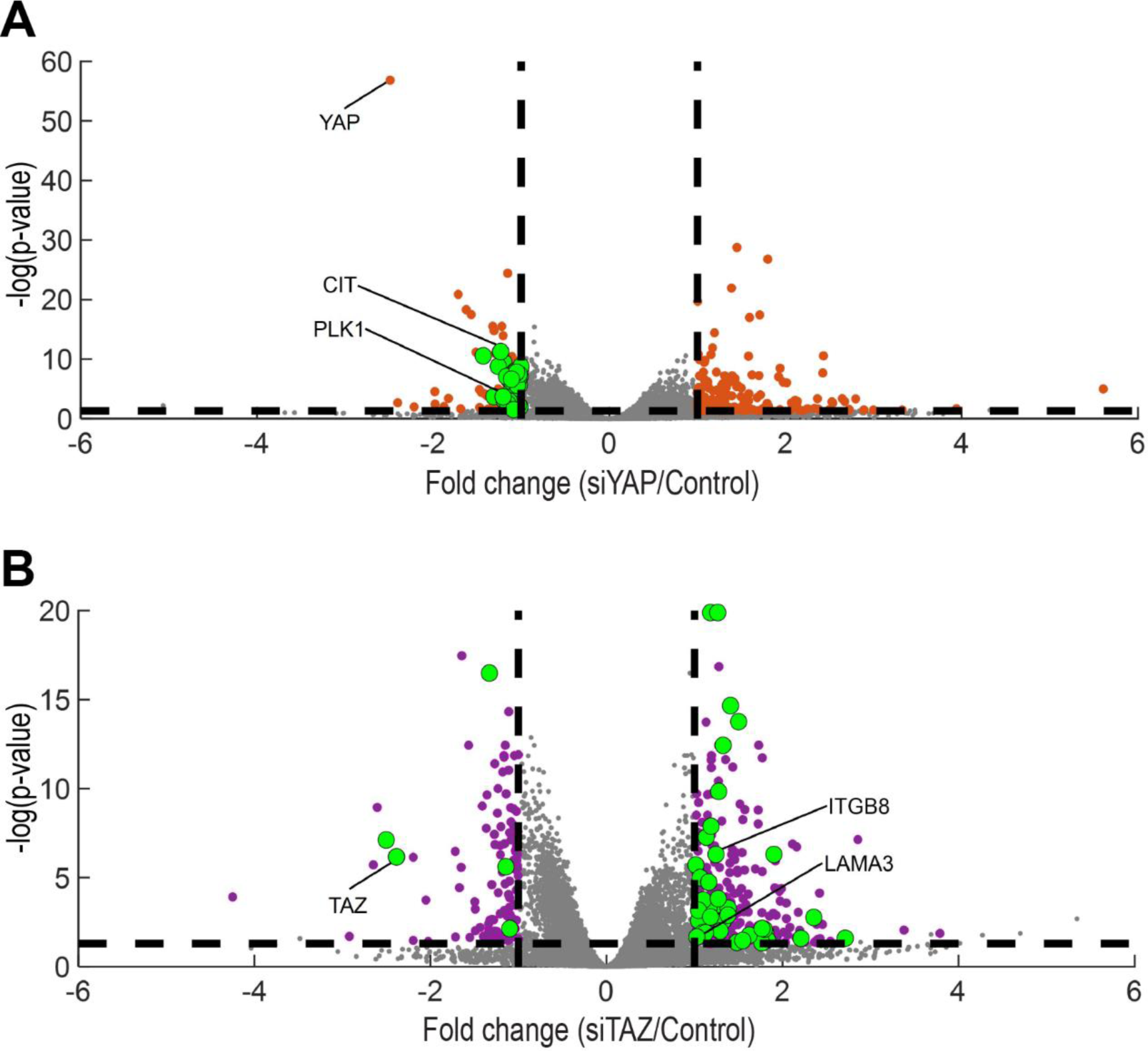
related to figure 2: Extended Volcano plots of YAP-regulated and TAZ-regulated genes. A and B. Volcano plots of all genes detected in the RNA-seq analysis. Colored dots represent genes whose expression was significantly altered by either siYAP (orange) or siTAZ (purple), relative to control siRNA. Genes related to the top enriched GO terms for each siRNA (black columns in Fig. 2A and 2B) are marked in green. Specific informative genes are indicated.

**Figure S3.**
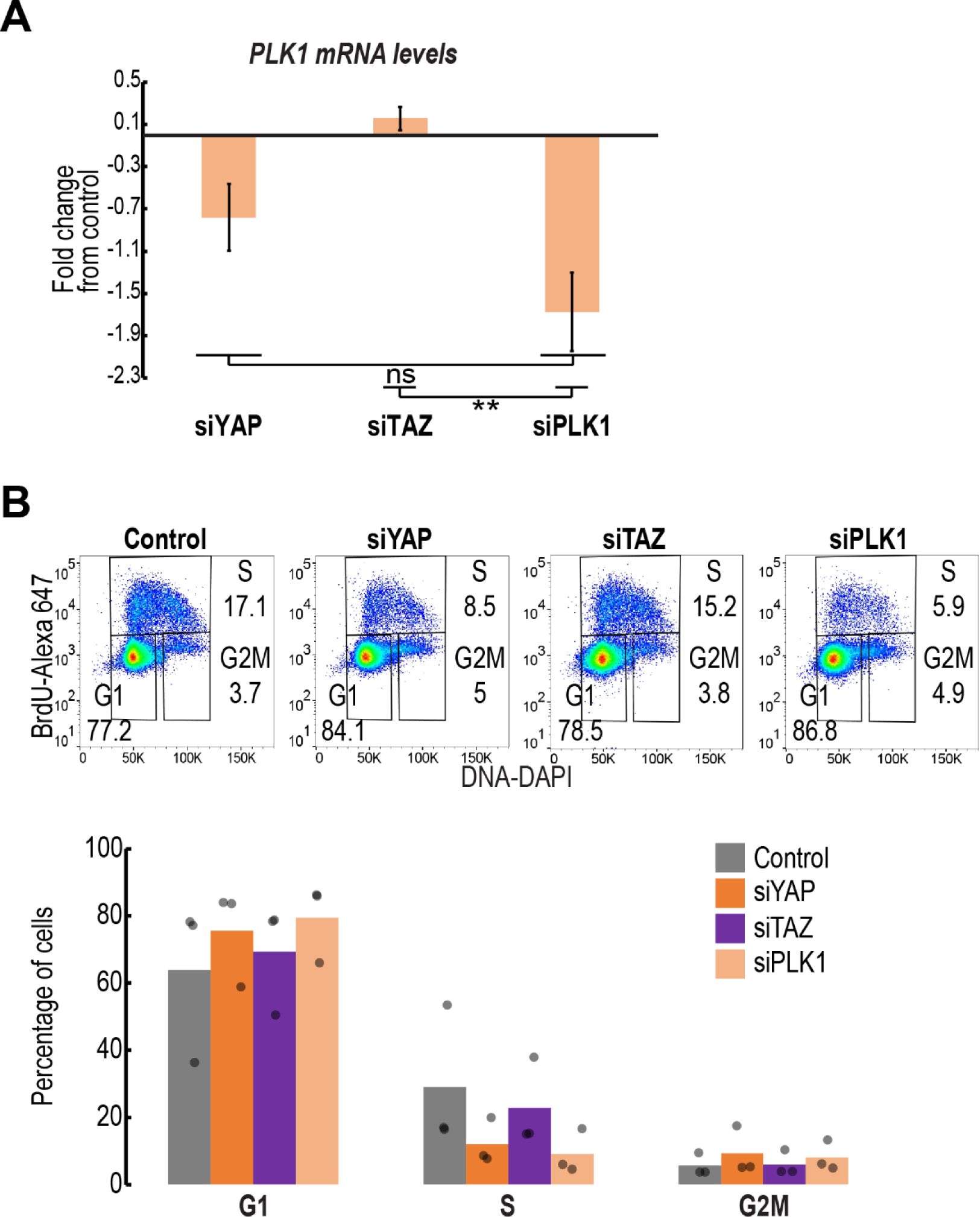
related to figure 3: silencing of PLK1 mimicked the cell cycle effects of siYAP. A. RT-qPCR analysis of PLK1 mRNA levels in H1299 cells transfected with the indicated siRNAs. Data represent log2 mRNA expression (mean+SEM) normalized to GAPDH and control transfected cells, from three independent biological repeats. ns = not significant; **p<0.01 one-way ANOVA and Tukey’s post hoc test of the indicated comparison. B. Cell cycle profiling by BrdU + DAPI of H1299 cells transfected with the indicated siRNAs, 48 hours post-transfection. Upper panel: representative FACS analysis images. Lower panel: Average percentages of cells in each cell cycle phase, from three biological repeats; each dot represents an independent biological repeat.

**Figure S4.**
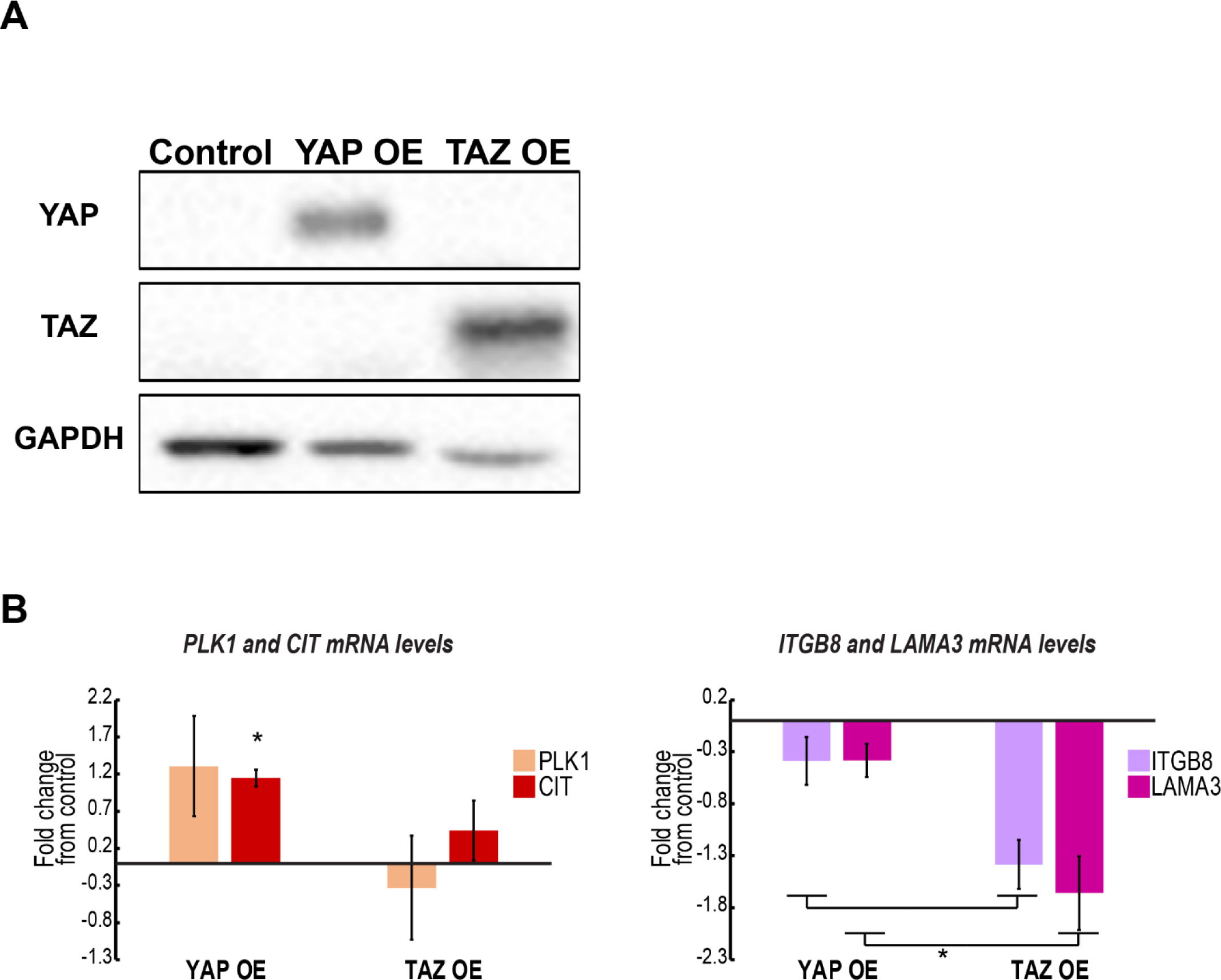
connected to figure 3 and figure 4: YAP and TAZ protein levels and candidate target gene expression upon YAP or TAZ overexpression. H1299 cells were transfected with plasmids encoding either YAP-flag (YAP OE) or TAZ-flag (TAZ OE). Cultures were harvested 48 hours post-transfection, including serum starvation in the last 18 hours. A. Representative Western blot analysis with antibodies specific to YAP, TAZ or GAPDH as loading control. B. RT-qPCR analysis of mRNA levels of the indicated genes. Data represent log2 mRNA expression (mean+SEM) normalized to GAPDH and control transfected cells, from three independent biological repeats. *p<0.05; one-way ANOVA and Tukey’s post hoc test for the indicated comparisons or vs. control.

**Figure S5.**
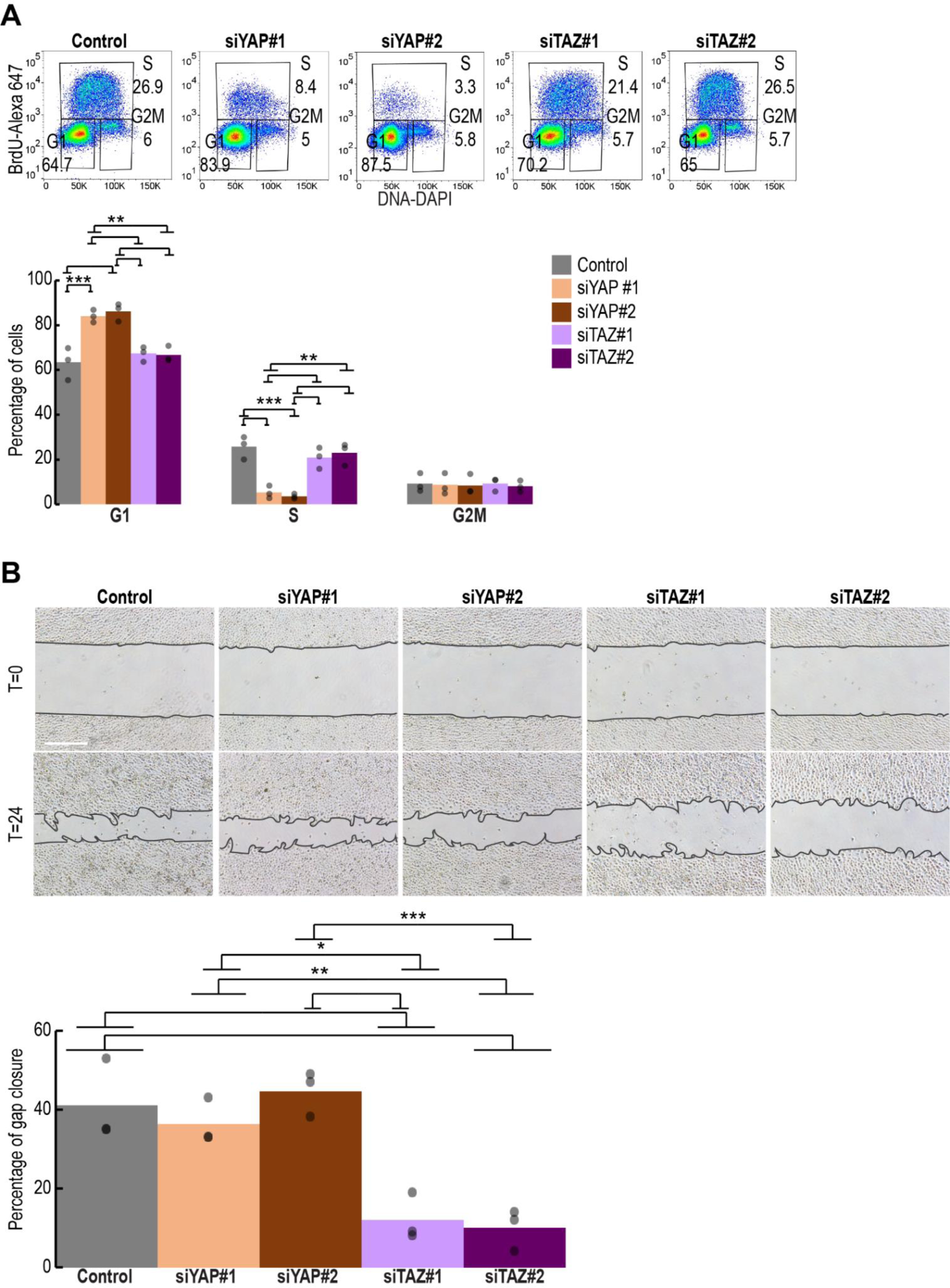
connected to figure 3 and figure 4: YAP or TAZ silencing by single siRNA oligonucleotides phenocopies the effects of YAP or TAZ silencing by siRNA SMARTpools. A. Cell cycle profiling by BrdU + DAPI of H1299 cell cultures (n=3) transfected with the indicated single siRNA oligonucleotides. Upper panel: representative FACS analysis images. Lower panel: Average percentages of cells in each cell cycle phase, from three biological repeats; each dot represents an independent biological repeat. **p<0.01; ***p<0.001 determined by one-way ANOVA and Tukey’s post hoc test of the indicated comparisons. B. Gap closure assay of H1299 cell cultures (n=3) transfected with the indicated siRNAs. Upper panel: representative images of gap closure at T=0 and T=24 hours. Lower panel: Average percentage of gap closure calculated from all biological repeats; each dot represents an independent biological repeat. *p<0.05; **p<0.01; ***p<0.001 one-way ANOVA and Tukey’s post hoc test of the indicated comparisons. Scale bar is 500µM.

**Figure S6.**
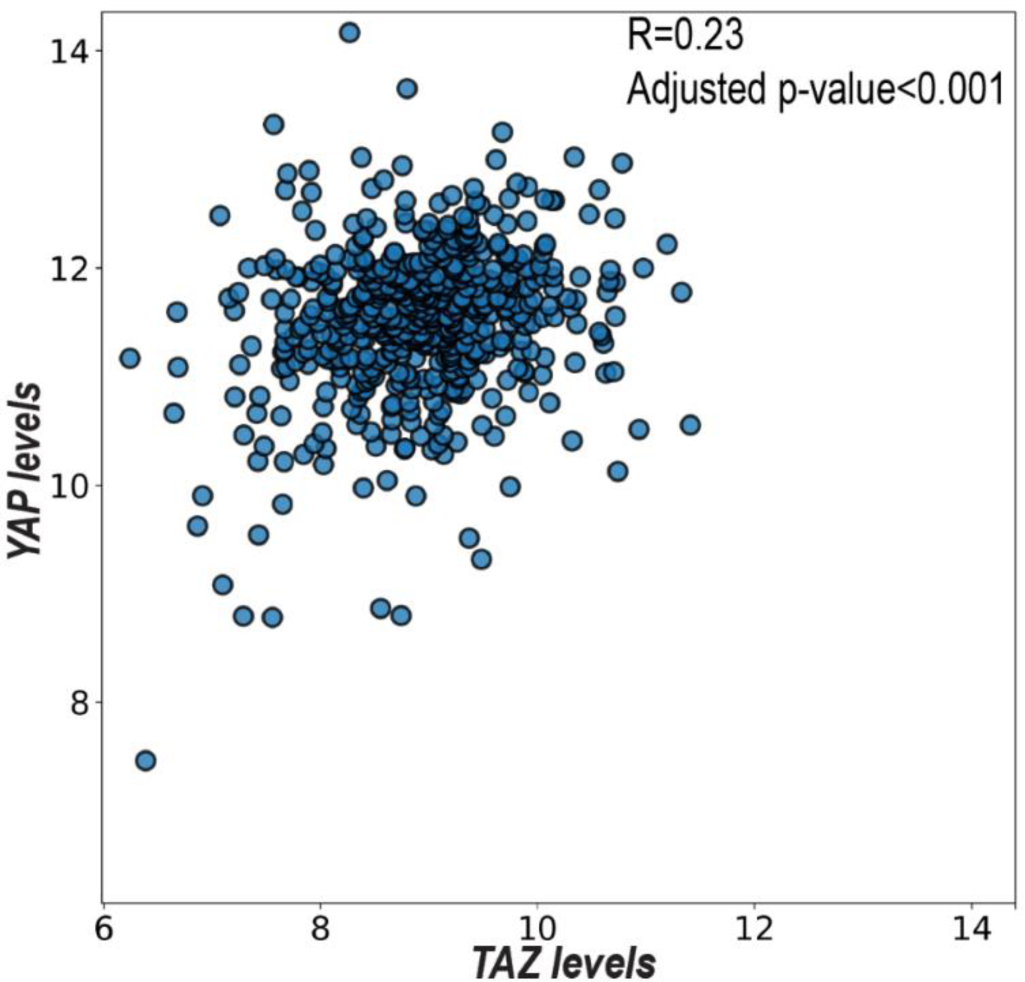
connected to figure 5: YAP and TAZ are only partially correlated across lung adenocarcinoma tumors. Scatter plot of YAP vs. TAZ mRNA levels across LUAD, derived from TCGA dataset. R = Pearson correlation coefficient. P-value was adjusted for multiple testing, using the procedure of Benjamini and Hochberg.

**Figure S7.**
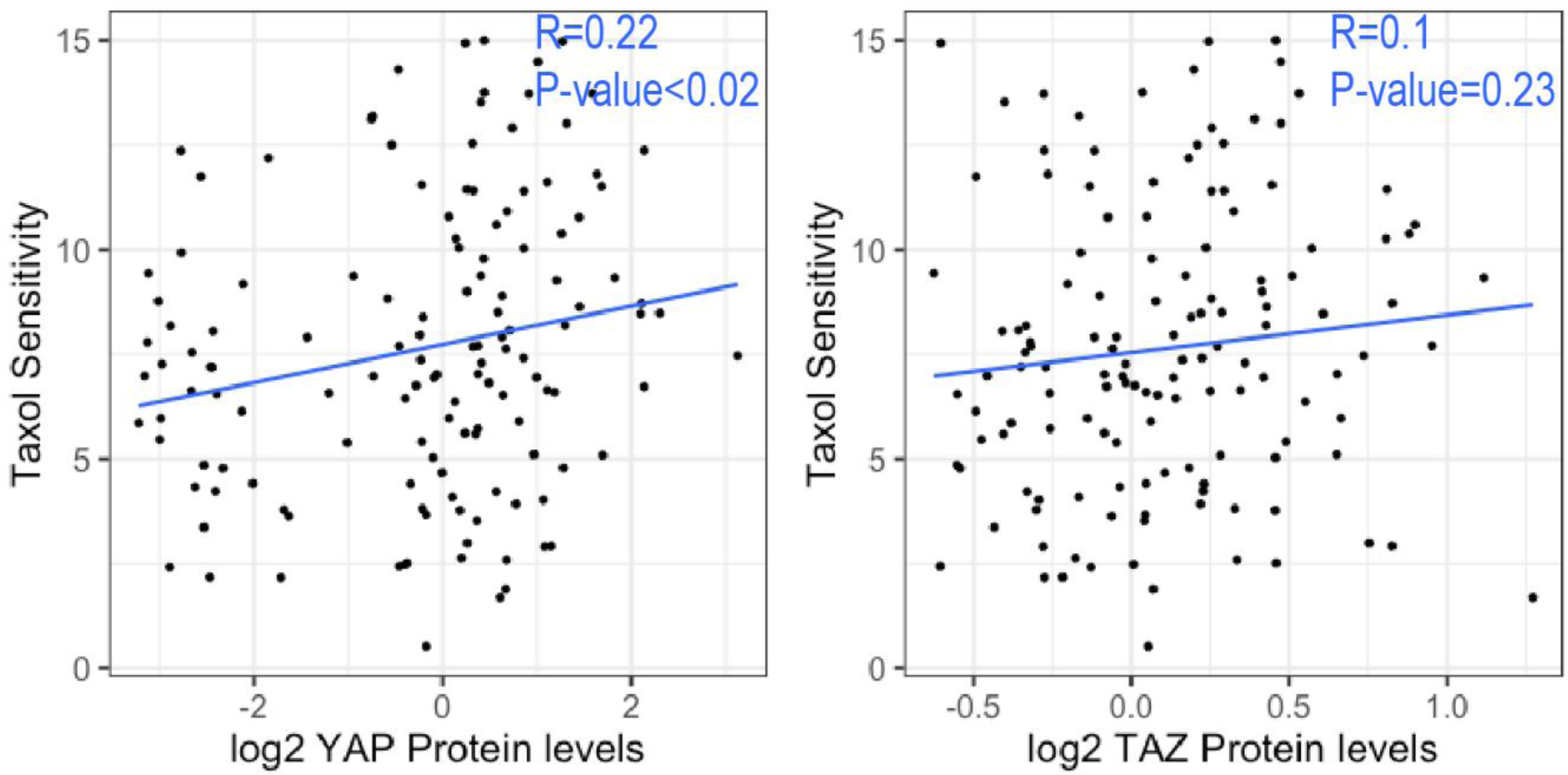
connected to figure 6: Taxol sensitivity is more correlated with YAP protein levels than with TAZ protein levels. Scatter plot of Taxol sensitivity in the GDSC dataset [40] (measured by area under the dose-response curve) plotted against YAP (left) or TAZ (right) protein levels (determined by RPPA in the CCLE) across lung cancer cell lines (n=136). Protein levels are log-transformed. R represents Spearman correlation coefficient, and blue line represents the indicated R value.

***Supplementary Table S1: RNA-seq Metascape Gene ontology results***

*Gene ontology results of either the YAP-regulated or TAZ-regulated genes from the RNA-seq analysis, determined by Metascape*.

***Supplementary Table S2: Primers and siRNA oligos***

*List of siRNA oligos and primers used in this study*.

***Supplementary Table S3: Table S3 -TCGA Metascape Gene ontology results***

*Gene ontology results of either the YAP-correlated or TAZ-correlated genes in LUAD TCGA, determined by Metascape*.

***Supplementary Table S4: List of YAP-differential and TAZ-differential drugs***

*List of 142 YAP-differential drugs and 38 TAZ-differential drugs in LUAD cells. Each row is a drug, and each column specifies additional information, such as: dose used in the screen, drug targets, indication, clinical phase and the p-value of the comparison*.

## References

1. Herbst, S.R., John V. Heymach, and S.M. Lippman, Lung cancer. The new england journal of medicine, 2008. 359(13): p. 1367–80.

2. Noguchi, S., et al., An integrative analysis of the tumorigenic role of TAZ in human non-small cell lung cancer. Clin Cancer Res, 2014. 20(17): p. 4660–72.

3. Wang, Y., et al., Overexpression of yes-associated protein contributes to progression and poor prognosis of non-small-cell lung cancer. Cancer Sci, 2010. 101(5): p. 1279–85.

4. Zhou, Z., et al., TAZ is a novel oncogene in non-small cell lung cancer. Oncogene, 2011. 30(18): p. 2181–6.

5. Lau, A.N., et al., Tumor-propagating cells and Yap/Taz activity contribute to lung tumor progression and metastasis. EMBO J, 2014. 33(5): p. 468–81.

6. Yeung, B., J. Yu, and X. Yang, Roles of the Hippo pathway in lung development and tumorigenesis. Int J Cancer, 2016. 138(3): p. 533–9.

7. Lo Sardo, F., S. Strano, and G. Blandino, YAP and TAZ in Lung Cancer: Oncogenic Role and Clinical Targeting. Cancers (Basel), 2018. 10(5).

8. Wang, Y., et al., Comprehensive Molecular Characterization of the Hippo Signaling Pathway in Cancer. Cell Rep, 2018. 25(5): p. 1304–1317.

9. Chen, H.Y., et al., R331W Missense Mutation of Oncogene YAP1 Is a Germline Risk Allele for Lung Adenocarcinoma With Medical Actionability. J Clin Oncol, 2015. 33(20): p. 2303–10.

10. Zanconato, F., et al., Genome-wide association between YAP/TAZ/TEAD and AP-1 at enhancers drives oncogenic growth. Nat Cell Biol, 2015. 17(9): p. 1218–27.

11. Beyer, T.A., et al., Switch enhancers interpret TGF-beta and Hippo signaling to control cell fate in human embryonic stem cells. Cell Rep, 2013. 5(6): p. 1611–24.

12. Hiemer, S.E., A.D. Szymaniak, and X. Varelas, The transcriptional regulators TAZ and YAP direct transforming growth factor beta-induced tumorigenic phenotypes in breast cancer cells. J Biol Chem, 2014. 289(19): p. 13461–74.

13. Galli, G.G., et al., YAP Drives Growth by Controlling Transcriptional Pause Release from Dynamic Enhancers. Mol Cell, 2015. 60(2): p. 328–37.

14. Callus, B.A., et al., YAPping about and not forgetting TAZ. FEBS Lett, 2018.

15. Piccolo, S., S. Dupont, and M. Cordenonsi, The biology of YAP-TAZ: Hippo signaling and beyond. Physiol Rev, 2014. 94: p. 1287–1312.

16. Yu, F.X. and K.L. Guan, The Hippo pathway: regulators and regulations. Genes Dev, 2013. 27(4): p. 355–71.

17. Elbediwy, A., et al., Integrin signalling regulates YAP and TAZ to control skin homeostasis. Development, 2016. 143(10): p. 1674–87.

18. Davis, J.R. and N. Tapon, Hippo signalling during development. Development, 2019. 146(18).

19. Finch-Edmondson, M. and M. Sudol, Framework to function: mechanosensitive regulators of gene transcription. Cell Mol Biol Lett, 2016. 21: p. 28.

20. Yuen, H.F., et al., TAZ expression as a prognostic indicator in colorectal cancer. PLoS One, 2013. 8(1): p. e54211.

21. Grijalva, J.L., et al., Dynamic alterations in Hippo signaling pathway and YAP activation during liver regeneration. Am J Physiol Gastrointest Liver Physiol, 2014. 307(2): p. G196–204.

22. Liu, C.Y., et al., MRTF/SRF dependent transcriptional regulation of TAZ in breast cancer cells Oncotarget, 2016.

23. Gao, Y., et al., YAP inhibits squamous transdifferentiation of Lkb1-deficient lung adenocarcinoma through ZEB2-dependent DNp63 repression. Nature Communications, 2014. 5.

24. Gobbi, G., et al., The Hippo pathway modulates resistance to BET proteins inhibitors in lung cancer cells. Oncogene, 2019. 38(42): p. 6801–6817.

25. Cohen, A.A., et al., Dynamic proteomics of individual cancer cells in response to a drug. Science, 2008. 322(5907): p. 1511–6.

26. Spolverini, A., et al., let-7b and let-7c microRNAs promote histone H2B ubiquitylation and inhibit cell migration by targeting multiple components of the H2B deubiquitylation machinery. Oncogene, 2017. 36(42): p. 5819–5828.

27. Furth, N., et al., LATS1 and LATS2 suppress breast cancer progression by maintaining cell identity and metabolic state. Life Sci Alliance, 2018. 1(5): p. e201800171.

28. Kohen, R., et al., UTAP: User-friendly Transcriptome Analysis Pipeline. BMC Bioinformatics, 2019. 20(1): p. 154.

29. Martin, M., Cutadapt removes adapter sequences from high-throughput sequencing reads. EMBnet.journal, 2011.

30. Dobin, A., et al., STAR: ultrafast universal RNA-seq aligner. Bioinformatics, 2013. 29(1): p. 15–21.

31. Love, M.I., W. Huber, and S. Anders, Moderated estimation of fold change and dispersion for RNA-seq data with DESeq2. Genome Biol, 2014. 15(12): p. 550.

32. Zhou, Y., et al., Metascape provides a biologist-oriented resource for the analysis of systems-level datasets. Nat Commun, 2019. 10(1): p. 1523.

33. Subramanian, A., et al., Gene set enrichment analysis: A knowledge-based approach for interpreting genome-wide expression profiles. PNAS, 2005. 102(43): p. 15545–15550.

34. Lee, J.S., et al., Harnessing synthetic lethality to predict the response to cancer treatment. Nat Commun, 2018. 9(1): p. 2546.

35. Viswanathan, S.R., et al., Genome-scale analysis identifies paralog lethality as a vulnerability of chromosome 1p loss in cancer. Nat Genet, 2018. 50(7): p. 937–943.

36. Weinstein, J.N., et al., The Cancer Genome Atlas Pan-Cancer analysis project. Nat Genet, 2013. 45(10): p. 1113–20.

37. Barretina, J., et al., The Cancer Cell Line Encyclopedia enables predictive modelling of anticancer drug sensitivity. Nature, 2012. 483(7391): p. 603–7.

38. Corsello, S.M., et al., Non-oncology drugs are a source of previously unappreciated anti-cancer activity. bioRxiv, 2019. doi.org/10.1101/730119.

39. Wishart, D.S., et al., DrugBank 5.0: a major update to the DrugBank database for 2018. Nucleic Acids Res, 2018. 46(D1): p. D1074–D1082.

40. Yang, W., et al., Genomics of Drug Sensitivity in Cancer (GDSC): a resource for therapeutic biomarker discovery in cancer cells. Nucleic Acids Res, 2013. 41(Database issue): p. D955–61.

41. Ghandi, M., et al., Next-generation characterization of the Cancer Cell Line Encyclopedia. Nature, 2019. 569(7757): p. 503–508.

42. Liu, J., et al., Genome and transcriptome sequencing of lung cancers reveal diverse mutational and splicing events. Genome Res, 2012. 22(12): p. 2315–27.

43. Tripathi, S.C., et al., Immunoproteasome deficiency is a feature of non-small cell lung cancer with a mesenchymal phenotype and is associated with a poor outcome. Proc Natl Acad Sci U S A, 2016. 113(11): p. E1555–64.

44. Lin, X.Y., et al., Expression of LATS1 contributes to good prognosis and can negatively regulate YAP oncoprotein in non-small-cell lung cancer. Tumour Biol, 2014. 35(7): p. 6435–43.

45. Ben-Porath, I., et al., An embryonic stem cell-like gene expression signature in poorly differentiated aggressive human tumors. Nat Genet, 2008. 40(5): p. 499–507.

46. Nardone, G., et al., YAP regulates cell mechanics by controlling focal adhesion assembly. Nat Commun, 2017. 8: p. 15321.

47. Wu, Z., et al., Up-regulation of CIT promotes the growth of colon cancer cells. Oncotarget, 2017. 8(42): p. 71954–71964.

48. Cappello, P., et al., Role of Nek2 on centrosome duplication and aneuploidy in breast cancer cells. Oncogene, 2014. 33(18): p. 2375–84.

49. Medina-Aguilar, R., et al., Resveratrol inhibits cell cycle progression by targeting Aurora kinase A and Polo-like kinase 1 in breast cancer cells. Oncol Rep, 2016. 35(6): p. 3696–704.

50. Hamill, K.J., A.S. Paller, and J.C. Jones, Adhesion and migration, the diverse functions of the laminin alpha3 subunit. Dermatol Clin, 2010. 28(1): p. 79–87.

51. Xu, Z. and R. Wu, Alteration in metastasis potential and gene expression in human lung cancer cell lines by ITGB8 silencing. Anat Rec (Hoboken), 2012. 295(9): p. 1446–54.

52. Giraldez, S., et al., G1/S phase progression is regulated by PLK1 degradation through the CDK1/betaTrCP axis. FASEB J, 2017. 31(7): p. 2925–2936.

53. Gupton, S.L. and C.M. Waterman-Storer, Spatiotemporal feedback between actomyosin and focal-adhesion systems optimizes rapid cell migration. Cell, 2006. 125(7): p. 1361–74.

54. Cereda, M., T.P. Mourikis, and F.D. Ciccarelli, Genetic Redundancy, Functional Compensation, and Cancer Vulnerability. Trends Cancer, 2016. 2(4): p. 160–162.

55. Helming, K.C., et al., ARID1B is a specific vulnerability in ARID1A-mutant cancers. Nat Med, 2014. 20(3): p. 251–4.

56. Horton, E.R., et al., Definition of a consensus integrin adhesome and its dynamics during adhesion complex assembly and disassembly. Nat Cell Biol, 2015. 17(12): p. 1577–1587.

57. Shreberk-Shaked, M. and M. Oren, New insights into YAP/TAZ nucleo-cytoplasmic shuttling: new cancer therapeutic opportunities? Mol Oncol, 2019. 13(6): p. 1335–1341.

58. Ramalingam, S. and C.P. Belani, Paclitaxel for non-small cell lung cancer. Expert Opin Pharmacother, 2004. 5(8): p. 1771–80.

59. Dandage, R. and C.R. Landry, Paralog dependency indirectly affects the robustness of human cells. Molecular Systems Biology, 2019. 15(9).

60. Sun, C., et al., Common and Distinctive Functions of the Hippo Effectors Taz and Yap in Skeletal Muscle Stem Cell Function. Stem Cells, 2017. 35(8): p. 1958–1972.

61. Plouffe, S.W., et al., The Hippo pathway effector proteins YAP and TAZ have both distinct and overlapping functions in the cell. J Biol Chem, 2018. 293(28): p. 11230–11240.

62. Tsai, C.R., et al., Yorkie regulates epidermal wound healing in Drosophila larvae independently of cell proliferation and apoptosis. Dev Biol, 2017. 427(1): p. 61–71.

63. Lin, T.H., et al., The Hippo pathway controls border cell migration through distinct mechanisms in outer border cells and polar cells of the Drosophila ovary. Genetics, 2014. 198(3): p. 1087–99.

64. Ren, F., et al., Hippo signaling regulates Drosophila intestine stem cell proliferation through multiple pathways. Proc Natl Acad Sci U S A, 2010. 107(49): p. 21064–9.

65. Huang, J., et al., The Hippo signaling pathway coordinately regulates cell proliferation and apoptosis by inactivating Yorkie, the Drosophila Homolog of YAP. Cell, 2005. 122(3): p. 421–34.

66. Thompson, B.J. and S.M. Cohen, The Hippo pathway regulates the bantam microRNA to control cell proliferation and apoptosis in Drosophila. Cell, 2006. 126(4): p. 767–74.

67. Morin-Kensicki, E.M., et al., Defects in yolk sac vasculogenesis, chorioallantoic fusion, and embryonic axis elongation in mice with targeted disruption of Yap65. Mol Cell Biol, 2006. 26(1): p. 77–87.

68. Hossain, Z., et al., Glomerulocystic kidney disease in mice with targeted inactivation of Wwtr1. Proceedings of the National Academy of Sciences of the United States of America, 2007. 104(5).

69. Makita, R., et al., Multiple renal cysts, urinary concentration defects, and pulmonary emphysematous changes in mice lacking TAZ. Am J Physiol Renal Physiol, 2008. 294(3): p. F542–53.

70. Wu, T., et al., Phase separation of TAZ compartmentalizes the transcription machinery to promote gene expression. bioRxiv, 2019. doi.org/10.1101/671230.

71. Cai, D., et al., Phase separation of YAP reorganizes genome topology for long-term YAP target gene expression. Nat Cell Biol, 2019. 21(12): p. 1578–1589.

